# Different growth pattern during microbial electrosynthesis using *C. ljungdahlii* evolutionary adapted on iron

**DOI:** 10.1101/2025.09.13.675759

**Authors:** Chaeho Im, Kaspar Valgepea, Oskar Modin, Yvonne Nygård, Carl Johan Franzén

## Abstract

*Clostridium ljungdahlii* is an acetogen used for syngas fermentation and capable of microbial electrosynthesis. While *C. ljungdahlii* has potential for industrial application because of its broad spectrum of metabolites and substrates, including CO_2_ and CO, the efficiency of extracellular electron uptake in a bioelectrochemical system is low. Therefore, *C. ljungdahlii* was evolutionary adapted on iron as sole electron source with the aim to improve extracellular electron uptake. Over 38 batches, 95% of the culture was replaced with fresh medium biweekly to retain iron-attached *C. ljungdahlii*, leading to improved acetate production rates with each cycle. Eight isolated strains were tested on fructose, H_2_ and CO_2_, and iron to screen evolved mutants with desired mutations. Compared to the wild-type, growth on fructose was similar and growth on H_2_ and CO_2_ was, surprisingly, worse, with only minor differences between isolates. The isolated mutants produced acetate at a rate of 0.14 ± 0.01 mM/d on iron, while the wild-type strain produced 0.75 ± 0.14 mM/d. Whole genome sequencing of isolated mutants revealed 16 mutations, of which seven mutations were found in all isolates. Mostly, membrane proteins were mutated. In a BES reactor, acetate production ceased after day 1. The optical density (OD) of isolate #8 stopped increasing after day 2, reaching 0.122 ± 0.005, followed by the production of formate and ethanol. The wild-type strain continued to grow until day 4, reaching an OD of 0.177 ± 0.003. These results may indicate that *C. ljungdahlii* slows down growth and produces ethanol as an energy reserve as an evolutionary strategy for survival in an electron-limited environment.

## 1. Introduction

Bioelectrochemical systems (BESs) utilize the interactions between microorganisms and insoluble solid electrodes to shift the intracellular redox balance either towards an oxidative state or to supply electrons to microorganisms for reductive reactions. These electrochemical interactions can realise metabolic reactions that cannot be achieved in conventional fermentation process environments and can lead to higher target product yields (1, 2). Bioelectrochemical interactions are also exploited in bioleaching, bioremediations of harmful chemicals like nitrate and sulphate, or CO_2_ fixation and commodity chemicals production using electricity (3).

Several extracellular electron transfer (EET) mechanisms of electron-donating bacteria (anodophilic bacteria) have been unveiled, including direct electron transfer, electron shuttle-mediated electron transfer, and long-range electron transfer using nanowires (4). However, other than hydrogen-mediated electron uptake, EET mechanisms of electron-accepting (cathodophilic) bacteria are still under investigation (5). Extracellular electron uptake (EEU) mechanisms of autotrophic methanogens and autotrophic acetogens are receiving increased attention as these self-regenerating biocatalysts can convert CO_2_ into commodity chemicals such as methane, fatty acids, and the corresponding alcohols employing *in-situ* electrolysis for delivery of reducing equivalents (6, 7). While several approaches have been introduced in order to investigate the mechanisms of the EEU, the conclusions reached so far are hypothetical and/or based on indirect observations (8).

Biological oxidation of zero-valent iron (ZVI) is a process that is very similar to uptake of electrons from a cathode in a BES. Iron acts as an insoluble electron donor in aqueous solution, similar to a cathode in a BES. The standard redox potential of ZVI oxidation is –0.47 V vs SHE (eq. 1) (9), which means it can drive cathodic reactions in anoxic aqueous solutions, without any help of an external power source and/or three electrode systems that are used for BES operation, with a proper electron acceptor, protons (eq. 2 and 3).

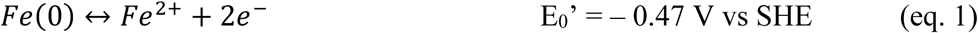

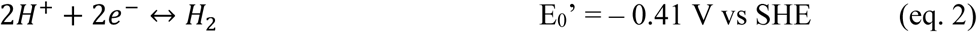

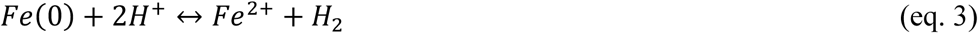

Interestingly, bacterial communities utilizing ZVI are very similar to bacterial communities in BES reactors (9–11). *Sporomusa* spp. were dominant during the first isolation of acetogens using iron. Similarly, *Acetobacterium* spp. emerged as the most dominant genus in two parallel experiments using iron as the sole electron source and CO_2_ as the carbon source. Furthermore, both *Sporomusa* spp. and *Acetobacterium* spp. were dominant when mixed cultures were cultivated under low hydrogen partial pressure (9, 11, 12). Similarly, *Acetobacterium* spp. were dominant when mixed culture was used in BESs in multiple independent studies (13–15). Also, several species from *Sporomusa* genus showed better performance in a BES than other homoacetogens (10, 16). Taken together, it is likely that the EEU linked to iron oxidation and to cathode oxidation share the same mechanism.

Adaptive laboratory evolution (ALE) is a method to investigate biological evolution by mimicking the natural environment of the microorganism or to improve the performance of a microorganism by enforcing selective pressure in a defined condition by cultivating them in serially in flasks or continuously in a bioreactor (17, 18). ALE for improving nutrient utilization and environmental stress tolerance is routinely done for developing industrial host organisms. *Clostridium tyrobutyricum* and *Eubacterium limosum* have been evolutionary adapted on their natural substrates, glucose and syngas, respectively. Evolved mutants showed significantly improved tolerance against high concentrations of each substrate and better productivity for metabolites (19, 20). This approach is also found in the microbial electrochemistry field. *Geobacterium sulfurreducens* can naturally reduce insoluble metal oxides using conductive nanowires, while oxidizing organic acids (21). Microbial fuel cell (MFC) utilize this dissimilatory metal reduction ability to reduce a solid electrode in a BES (22). Therefore, *G. sulfurreducens* was cultivated long-term under microbial fuel cell (MFC) conditions and in serum bottles with the insoluble electron acceptor Fe(III) oxide to improve dissimilatory electrode reduction for electricity generation (22, 23). Evolved mutant strains isolated from each study showed superior abilities to reduce the anode in the MFC and the Fe(III) oxide, respectively (22, 23). However, long-term cultivation of *G. sulfurreducens* in a MFC required continuous flow of fresh medium in order to supply nutrients to support growth (23). On the other hand, *Sporomusa ovata* was adapted on methanol to accelerate autotrophic metabolism for improved performance of microbial electrosynthesis (24). *S. ovata* was serially cultivated in serum bottles containing 2% methanol as carbon and electron source, and acetate production rate was improved 6.5-fold. However, this approach can be applied only to homoacetogens equipped with the methanol assimilation pathway. Additionally, growth of pure culture homoacetogens in BESs is very low (25, 26). Consequently, it is difficult to cultivate homoacetogens in BESs for long enough to accumulate beneficial mutations in the electron uptake systems.

*Clostridium ljungdahlii*, one of the type-strains for syngas fermentation, is capable of microbial electrosynthesis in a BES (10). It has a broad natural product spectrum, producing formate, acetate, ethanol, lactate, and 2,3-butanediol through CO and CO_2_ fixation via the Wood-Ljungdahl Pathway. However, the ability of *C. ljungdahlii* to utilize extracellular electron from a cathode in a BES is moderate compared to other acetogens investigated so far (6, 10, 27). In a previous study, *C. ljungdahlii* demonstrated the capability for microbial electrosynthesis, whereas *A. woodii*, despite having a higher H_2_ affinity than *C. ljungdahlii*, was unable to utilize electrons from the electrode under the same conditions. This suggests the presence of an unknown electron uptake mechanism in *C. ljungdahlii* (10). While *S. ovata* can utilize electrons from a cathode in a BES with high efficiency (≥ 75 %) at various cathodic potentials, the coulombic efficiency of *C. ljungdahlii* in a BES decreased with more negative cathodic potentials (25, 28). Therefore, identification and improvement of *C. ljungdahlii*’s EEU in a BES reactor are required for its use as a cell factory in a BES.

In this study, a novel method to improve EEU of *C. ljungdahlii* using iron(0) was tested. *C. ljungdahlii* was adapted gradually on decreasing partial pressures of hydrogen until ZVI was used as the sole electron source. After 13 months of evolutionary adaptation (EA) in flasks with ZVI granules as the sole electron donor and continuous CO_2_ sparing as the sole carbon source, the evolved mutants were isolated. Isolated mutants were tested in different growth conditions and were genome sequenced to elaborate the performances compared to the wild-type. At the end, one of isolated mutants showed a different growth pattern in a BES.

## 2. Materials and Methods

### 2.1 Bacteria and culture conditions

*Clostridium ljungdahlii* (DSM-13528) was obtained from DSMZ (Braunschweig, Germany). The bacterial culture was stored as 1.5 mL aliquots in anoxic 20% glycerol at -80°C. The cryostock of the strain was streaked onto an YTF agar plate in an anaerobic chamber (Don Whitley Scientific Ltd, UK). Once colonies were visible, a single colony was inoculated in 40 mL of modified DSMZ 879 medium with and without 1 g/L of yeast extract under [H_2_:CO_2_] = [80:20] of 2 bars over-pressure for pre-cultivation in a 200 mL serum bottle. The modified DSMZ 879 medium contained (per liter): 2-ethanesulfonic acid monohydrate 30 g; NH_4_Cl 1g; KCl 0.1 g; MgSO_4_×7H_2_O 0.2 g; NaCl 0.8 g; KH_2_PO_4_ 0.01 g; CaCl_2_×2H_2_O 0.02 g; Sodium acetate 0.25 g; FeSO_4_×7H_2_O 50 mg; L-Cysteine×HCl 0.6 g; 0.1 % w/v Na-resazurin solution 1 mL; 10 mL of modified Wolin’s mineral solution (contents per liter: nitrilotriacetic acid 1.5 g; MgSO_4_×7H_2_O 3 g; MnSO_4_×H_2_O 0.5 g; NaCl 1 g; FeSO_4_×7H_2_O 0.1 g; CoSO_4_×7H_2_O 0.18 g; CaCl_2_×2H_2_O 0.1 g; ZnSO_4_×7H_2_O 0.18 g; CuSO_4_×5H_2_O 0.01 g; KAl(SO_4_)_2_×12H_2_O 0.02; H_3_BO_3_ 0.01 g; Na_2_MoO_4_×2H_2_O 0.01 g; NiCl_2_×6H_2_O 0.03 g; Na_2_SeO_3_×5H_2_O 0.3 mg; Na_2_WO_4_×2H_2_O 0.4 mg) and 10 mL of Wolin’s vitamin solution (contents per liter: biotin 2 mg; folic acid 2 mg; pyridoxine×HCl 10 mg; thiamine×HCl 5 mg; riboflavin 5 mg; nicotinic acid 5 mg; Ca×D-pantothenate 5 mg; vitamin B12 0.1 mg; p-aminobenzoic acid 5 mg; a-lipoic acid 5 mg.

### 2.2 Evolutionary adaptation of *C. ljungdahlii* on iron

Adaptive laboratory evolution on iron was done in two steps: 1) gradual adaptation of *C. ljungdahlii* on iron with decreasing amount of hydrogen available in the headspace; 2) repeated cultivation of *C. ljungdahlii* with iron as sole electron source and continuous CO_2_ sparging as carbon source.

*C. ljungdahlii* was cultivated initially in a serum bottle containing 2 g of iron granules filled with [H_2_:CO_2_] = [80:20] at 2 bars overpressure. When the optical density (OD) of the culture reached 0.1, 90 % of the culture was replaced with fresh medium, and the headspace was filled with [N_2_:H_2_:CO_2_]=[27:53:20] at 2 bars overpressure (Fig. 1).

**Figure 1.**
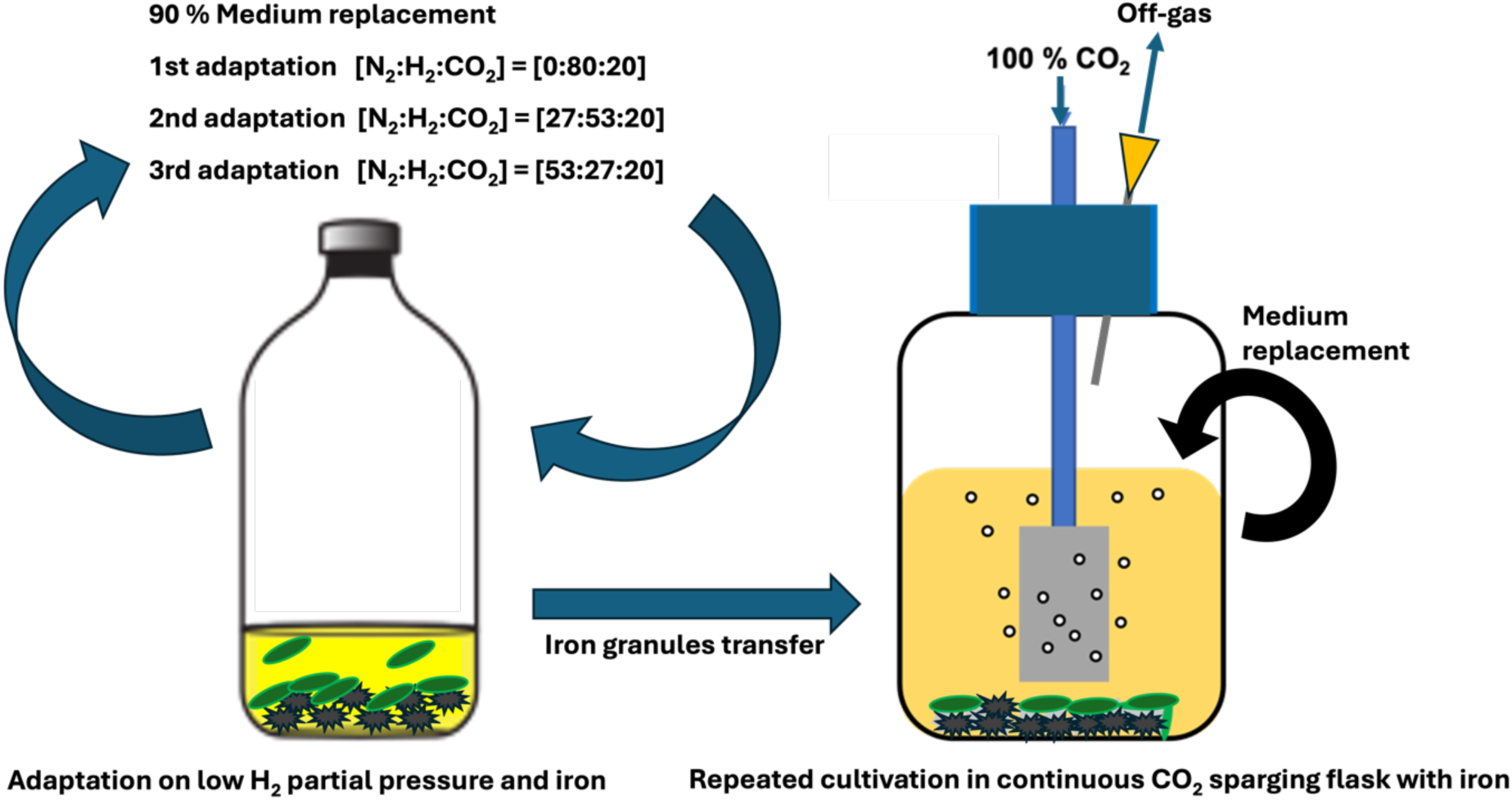
Schematic diagram of setup for evolutionary adaptation of *C. ljungdahlii* on iron

This process was repeated when the OD of the culture reached 0.1, with the headspace filled with a gas mixture of [N_2_:H_2_:CO_2_] = [53:27:20] at 2 bars overpressure. It was then assumed that *C. ljungdahlii* had adapted to low hydrogen partial pressure and the use of iron as an electron donor. Finally, 10 % of culture medium and the Fe^0^ granules in the culture medium were added to the bottles for long-term adaptation on iron with continuous 100 % CO_2_ sparging to remove hydrogen accumulated in the headspace. Each cultivation was maintained for 1.5 – 2 weeks until the pH of the culture increased to 5.7. At this point, 95 % of the culture was replaced with fresh growth medium.

When the performance of isolated mutants was tested, iron sheets having same surface area (3 cm x 2.5 cm) were used for the experiment in order to perform fair and reproducible comparisons of the isolates when grown on iron.

### 2.3 Isolation, selection, and storage of mutants to be tested

At the end of the long-term adaptation on iron, several iron granules were inoculated in a serum bottle filled with [H_2_:CO_2_] = [80:20] in the headspace to enrich evolved mutants. After 2 days of cultivation, the planktonic enrichment in the serum bottle was used for roll-tube method to be able to pick single colonies (29). When growth of single colonies was visible, 17 colonies were streaked on YTF agar plates to compare the morphologies and the growth of each mutant. At the same time, each colony was cultivated in YTF medium, and the cultures were stored and frozen in 20 % glycerol at -80°C until further use.

### 2.4 Bioelectrochemical system operation

A H-type BES reactor from Adams & Chittenden Scientific Glass Coop (Berkeley, US) with cathode and anode chambers connected by NW40 high temperature chain clamps (Grabs, Switzerland) and separated by a Nafion 117 membrane (Thermo Scientific Alfa Aesar) was used for the BES experiments. The BES contained 250 mL of modified DSMZ 879 medium omitting resazurin in each chamber. Graphite blocks (2.5 cm X 1.0 cm X 2.5 cm) connected with titanium wire (Alfa Aesar) were used as the cathode. Platinized titanium wire was used as the anode. 100 % of CO_2_ was sparged continuously at 10 mL/min to the BES cultures. For the BES experiments, 1.0 g/L of yeast extract was added to increase reproducibility among replicate experiments for the comparison between wild-type and the mutant (25). 10 mA of constant current was applied to a BES reactor by using a KP 07 potentiostat/galvanostat (Bank Elektronik - Intelligent Controls GmbH, Pohlheim, Germany) to further increase reproducibility without current fluctuation that sometimes occurs even when a constant cathodic potential is applied. An Ag/AgCl (3M NaCl) reference electrode (BASi, USA) was used to control the cathodic potential.

### 2.5 Analytical methods

During the experiments, samples were taken every 24 hours and the OD was measured at 660 nm using a spectrophotometer, WPA S1200+ Visible spectrophotometer (Biochrom™, UK). Metabolites analysis was performed using a high-performance liquid chromatography (HPLC) system equipped with a RI detector (40 °C) and a UV detector (210 nm) (Jasco, Japan). A ROA-Organic acid H+ (8%) column (Rezex, USA) was used to separate the metabolites, and 5 mM of H_2_SO_4_ eluent solution was pumped through the column at 0.6 mL/min. The oven temperature was maintained at 60 °C. The pH of the cathode chamber was measured with a pH meter (Mettler Toledo, USA).

### 2.6 Whole genome sequencing

Isolated mutants were cultivated in YTF medium overnight. 1 mL of overnight cultures were harvested in 1.5 mL eppendorf tubes and centrifuged at 12,000 RPM for 2 minutes. Next, genomic DNAs were extracted using MasterPure Gram Positive DNA purification kits (LGC Bioresearch Technologies, Hoddesdon, UK) according to the manufacturer’s protocol. The qualities and quantities of gDNA samples were evaluated using a NanoDrop 2000 spectrophotometer (Thermo Fisher Scientific, Germany). gDNA samples were sent to Macrogen Europe (Amsterdam, Netherlands) for whole genome sequencing. The sample libraries were prepared using TruSeq Nano DNA Library Prep kits according to TruSeq Nano DNA Sample Preparation Guide, Part #15041110 Rev. D. Sequencing was performed using a NovaSeq 6000 150 bp paired-end sequencing system (Illumina, California, US).

After sequencing, FastQC (v0.11.8) was performed in order to check the read quality and Trimmomatic (v0.38) was used to remove adapter sequences and low quality reads in order to reduce biases in analysis (30, 31). In each sample, filtered data were mapped using BWA (v0.7.17) with mem algorithm to the *C. ljungdahlii* reference genome (GeneBank ID: GCA_000143685.1) in NCBI RefSeq database (32). After read mapping, duplicated reads were removed with Sambamba (v0.6.8) (33). Genome coverage and mapping ratio of mapped reads on the reference genome were calculated and variant calling was performed using SAMtools (v.1.12) and BCFtools (v.1.12) (30, 34). In this step, SNPs and short indel candidates with phred score over 30 (base call accuracy of 99.9%) were captured using the information of the mapped reads. These variants were then classified by chromosomes or scaffolds, and the information of the location was marked. Captured variants were annotated with SnpEff (v.4.3t) in order to interpret the effects of genetic variants (35).

## 3. Results

### 3.1 Comparison of granular iron and iron powder for adaptation of *C. ljungdahlii*

In order to let *C. ljungdahlii* grow on metallic iron, it had to be gradually adapted onto ZVI as sole electron source by using decreasing amount of hydrogen available in the headspace with CO_2_ as carbon source. Initially, ZVI powder was added to the serum flasks filled with [H_2_:CO_2_]=[80:20] at 2 bars overpressure. As soon as the sterile growth medium was added to the serum flasks, the pH of the growth medium increased from pH 5.0 to 5.4, and after 2 days the pH of the growth medium had reached 6.2 (Fig. 2). That drastic increase of pH beyond optimal growth pH range (pH 4.0 – 6.0) of *C. ljungdahlii* could inhibit the cell growth (36).

**Figure 2.**
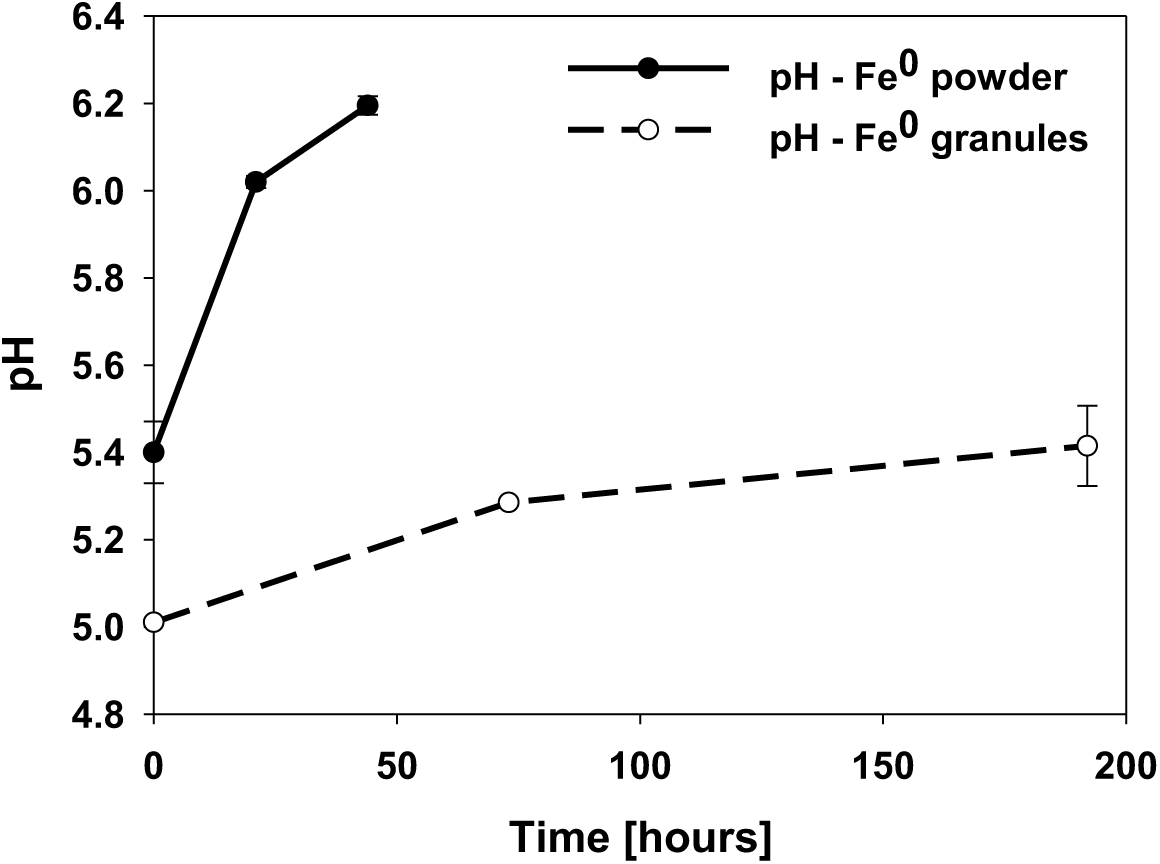
Media pH increase caused by Fe^0^ powder and granules oxidation. The error bars represent the standard deviations of the mean (n=2).

The pH increase is due to the ZVI corrosion reaction in anoxic condition (Eq 1.) (12). Here, when one mole ZVI is oxidized (acts as an electron donor), two moles of protons are required. Eventually, that causes an increase in the pH of the medium.

To slow down the pH increase, the ZVI powder was replaced by ZVI granules. This way the iron oxidation rate was reduced as the surface area was decreased. The pH increase became slower with ZVI granules than with ZVI powder, and the pH of the growth medium was well maintained below pH 5.5 for 192 hours (Fig. 2). Therefore, ZVI granules were chosen to be used for AE experiment.

### 3.2 Long-term adaptation of *C. ljungdahlii* on ZVI as sole electron source

After three serial transfers of *C. ljungdahlii* to ZVI under conditions with gradually decreasing hydrogen levels in the headspace, it was assumed that the *C. ljungdahlii* culture had adapted to using iron as the electron donor under low hydrogen partial pressure condition. 10% of the adapted culture along with ZVI granules in the serum bottle was inoculated in a flask containing 2 g of ZVI granules and was sparged with pure CO_2_ (see Materials and Methods). Acetate production of was detected after 4 days of cultivation with iron as the sole electron source. The acetate production rate was measured at the end of each batch. After the first cultivation in two parallel experiments, acetate was produced at the rate of 0.20 and 0.38 mM/d, respectively (Figure 3).

**Figure 3.**
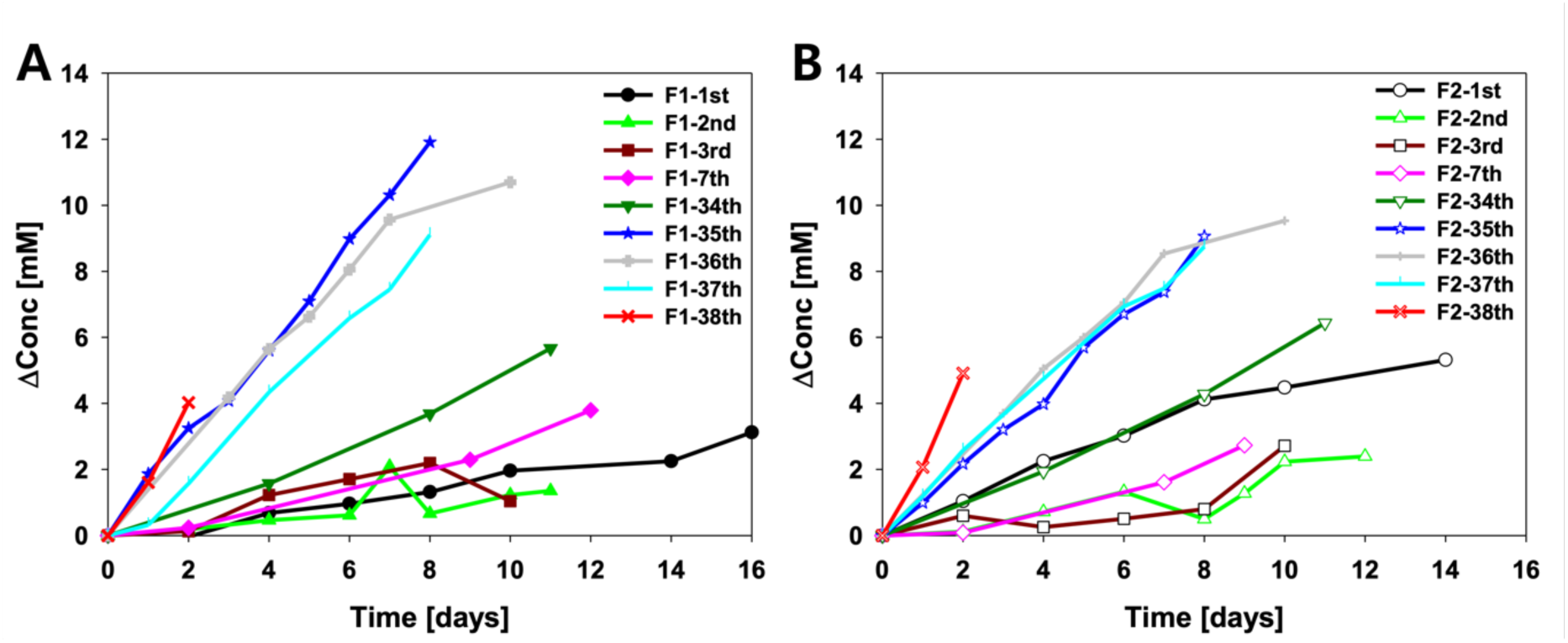
Acetate production profiles after the 1^st^ to 38^th^ serial medium replacement of two parallel experiments, F1 (A) and F2 (B). Flasks with iron granules were continuously sparged with CO_2_ over the evolutionary adaptation.

Since most planktonic cells were removed by medium replacement, the acetate production rate of each batch was decreased or remained around same level until the 7^th^ cultivation. Nonetheless, in the 34^th^ cultivation for F1 and in the 35^th^ cultivation for F2, the acetate production rate was doubled compared to the initial cultures, after which it continued to increase with each transfer (Figure 3). It was not possible to measure the growth of the cells due to color change of the media, formation of precipitates (e.g. Fe(OH)_2_, Fe(OH)_3_, and Fe_2_(CO_3_)_3_), and attachment of cells on the iron granule surface (e.g. cells growing on iron surface, which was the primary target of the experiment). The acetate production rate in the two series F1 and F2 increased by 10-fold and 6.5-fold, respectively, in the 38th cultivation compared to the initial cultivations.

### 3.3 Screening for an evolved strain performing better in a BES

A total of 17 isolates were picked after the ALE experiment. Based on different morphologies and growth of colonies on agar plates, 8 isolates were tested for changes in growth on fructose or H_2_:CO_2_. The specific growth rate on fructose of all the isolates was 5 – 15 % higher compared to the wild-type strain (Figure 4).

**Figure 4.**
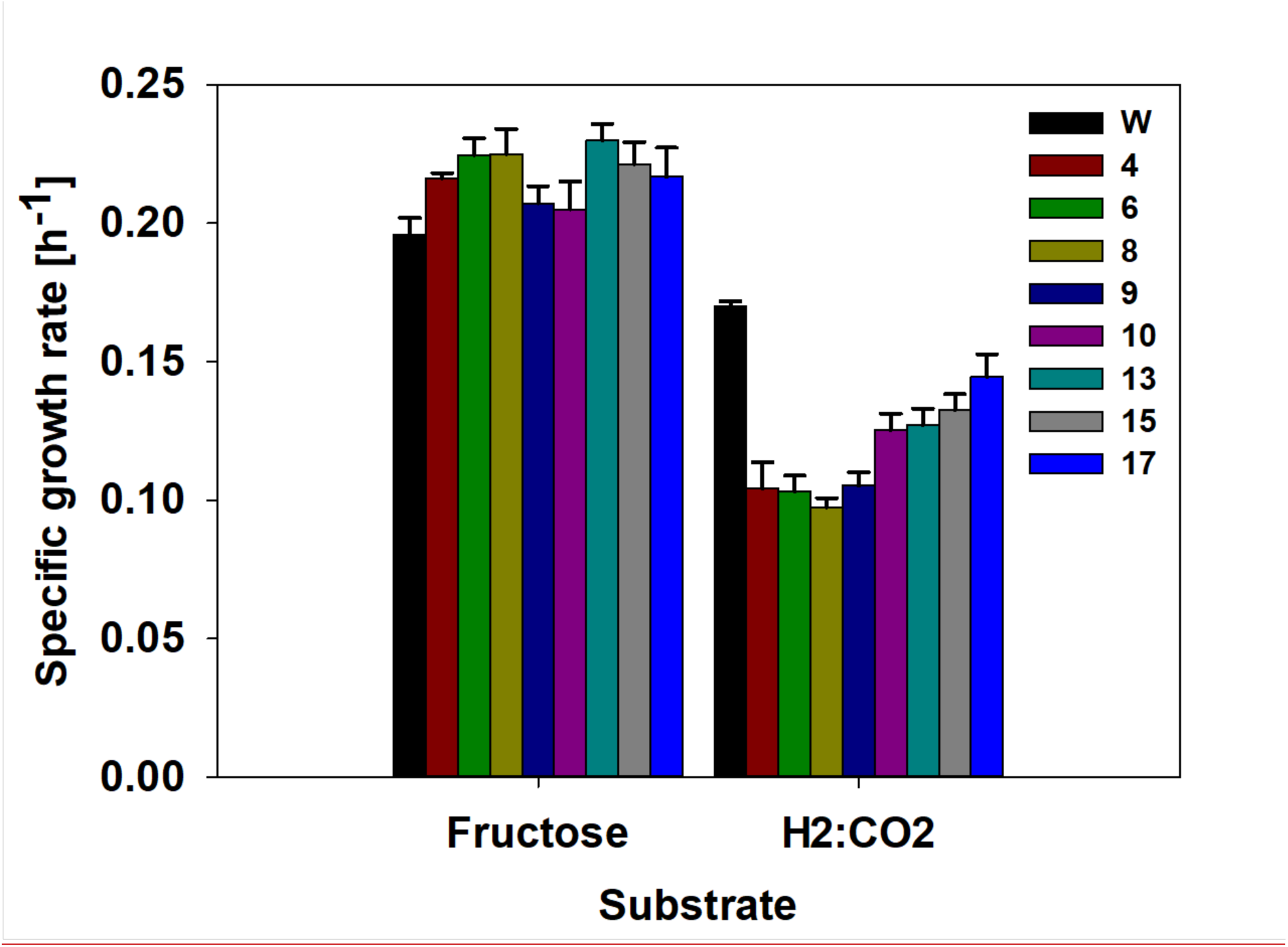
Specific growth rate of wild-type (W) and selected isolates on fructose and H_2_:CO_2_. The error bars represent standard deviations of the mean (n=3).

On the other hand, all of the isolates grew more slowly on H_2_/CO_2_ compared to the wild-type strain. There was no difference in metabolite production of the strains grown on H_2_:CO_2_.

### 3.4 Performance of evolved mutants on iron as electron source

Five isolates with either fast (#17), average (#10), or slow (#4, #6, and #8) autotrophic growth were selected for further evaluation. The wild-type strain outperformed the isolates with an acetate production rate of 0.75 ± 0.14 mM/d under CO_2_ as a carbon source and iron as a sole electron donor. The other isolates produced acetate at less than 0.14 ± 0.01 mM of acetate /d (Figure 5).

**Figure 5.**
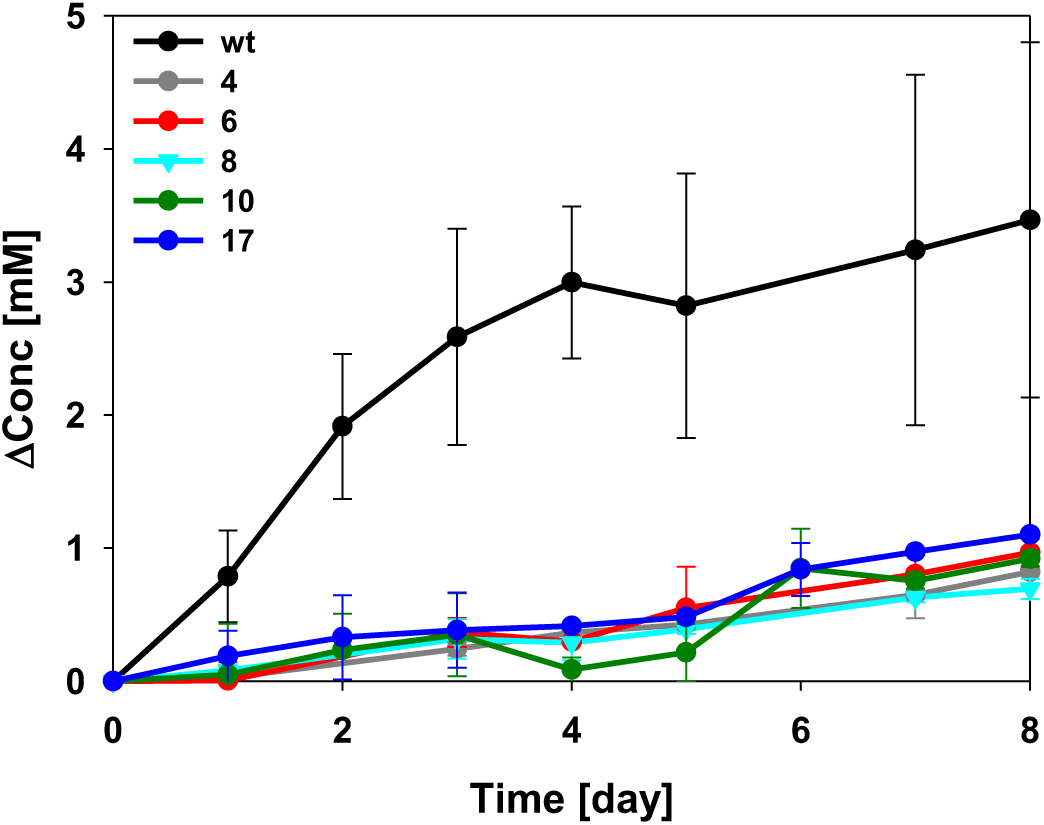
Acetate production profile of *C. ljungdahlii* wild-type and selected isolates when grown on CO_2_ as a carbon source and iron as a sole electron source. The error bars represent the standard deviations of the mean (n=2).

### 3.5 Detected mutations in the isolates

Whole-genome sequencing of the isolated mutants was performed to identify mutations that occurred during evolutionary adaptation on iron and to investigate the performance of the isolated mutants on iron compared to the wild-type strain. The genome of the parental strain used for ALE was sequenced to identify the mutations that happened during reactivation of the freeze-dried culture, freeze-thaw cycle for storage at - 80 °C, and propagation and to avoid mis-interpretation of the results from whole genome sequencing of isolated mutants (Table 1).

**Table 1.** Mutations detected in the wild-type (W) parental strain for the evolutionary adaptation compared to the reference genome.

| Position (bp) | Mutation | Amino acid change | Gene ID | Annotation |
| --- | --- | --- | --- | --- |
| 476,961 | A → G | Y350C | CLJU_RS02190 | Glutamyl-tRNA reductase |
| 683,222 | ATTT → ATTTT | intergenic | CLJU_RS03040 | Type 1 glutamine amidotransferase |
| 1,400,774 | G → T | S227I | CLJU_RS06385 | GTP-sensing pleiotropic transcriptional regulator |
| 1,614,559 | G → A | Intergenic | CLJU_RS07295 | Helix-turn-helix transcriptional regulator |
| 1,614,561 | C → T | Intergenic | CLJU_RS07295 | Helix-turn-helix transcriptional regulator |
| 1,619,490 | CT → C | intergenic | CLJU_RS07330 | Hypothetical protein |
| 1,817,665 | T → C | Intergenic | CLJU_RS08235 | Hypothetical protein |
| 2,235,661 | T → G | I129S | CLJU_RS10055 | Biotin-dependent carboxyltransferase family protein:<br>5-oxoprolinase (ATP-hydrolysing) |
| 2,595,040 | A → T | Synonymous<br>A355A | CLJU_RS11595 | Uracil permease |
| 2,698,277 | TAAA → TAAAA | frameshift | CLJU_RS22935 | Hypothetical protein |
| 3,211,628 | A → AAATATGTT | intergenic | CLJU_RS14330 | Fructose-bisphosphatase class III |
| 3,352,599 | C → T | Synonymous<br>V785V | CLJU_RS14955 | Lamin tail domain-containing protein<br>(ion channels) |
| 3,352,602 | T → C | Synonymous<br>T784T | CLJU_RS14955 |  |
| 3,352,611 | C → T | Synonymous<br>G781G | CLJU_RS14955 |  |
| 3,352,677 | C → T | Synonymous<br>T759T | CLJU_RS14955 |  |
| 3,352,736 | G → A | Synonymous<br>L740L | CLJU_RS14955 |  |
| 3,352,780 | T → G | E725A | CLJU_RS14955 |  |
| 3,352,950 | T → C | Synonymous<br>T668T | CLJU_RS14955 |  |
| 3,352,953 | T → C | Synonymous<br>V667V | CLJU_RS14955 |  |
| 3,352,956 | C → T | Synonymous<br>V666V | CLJU_RS14955 |  |
| 3,353,396 | C → T | V520I | CLJU_RS14955 |  |
| 4,428,830 | A → C | I161M | CLJU_RS20060 | Energy coupling factor transporter transmembrane<br>protein |

**Table 3.**
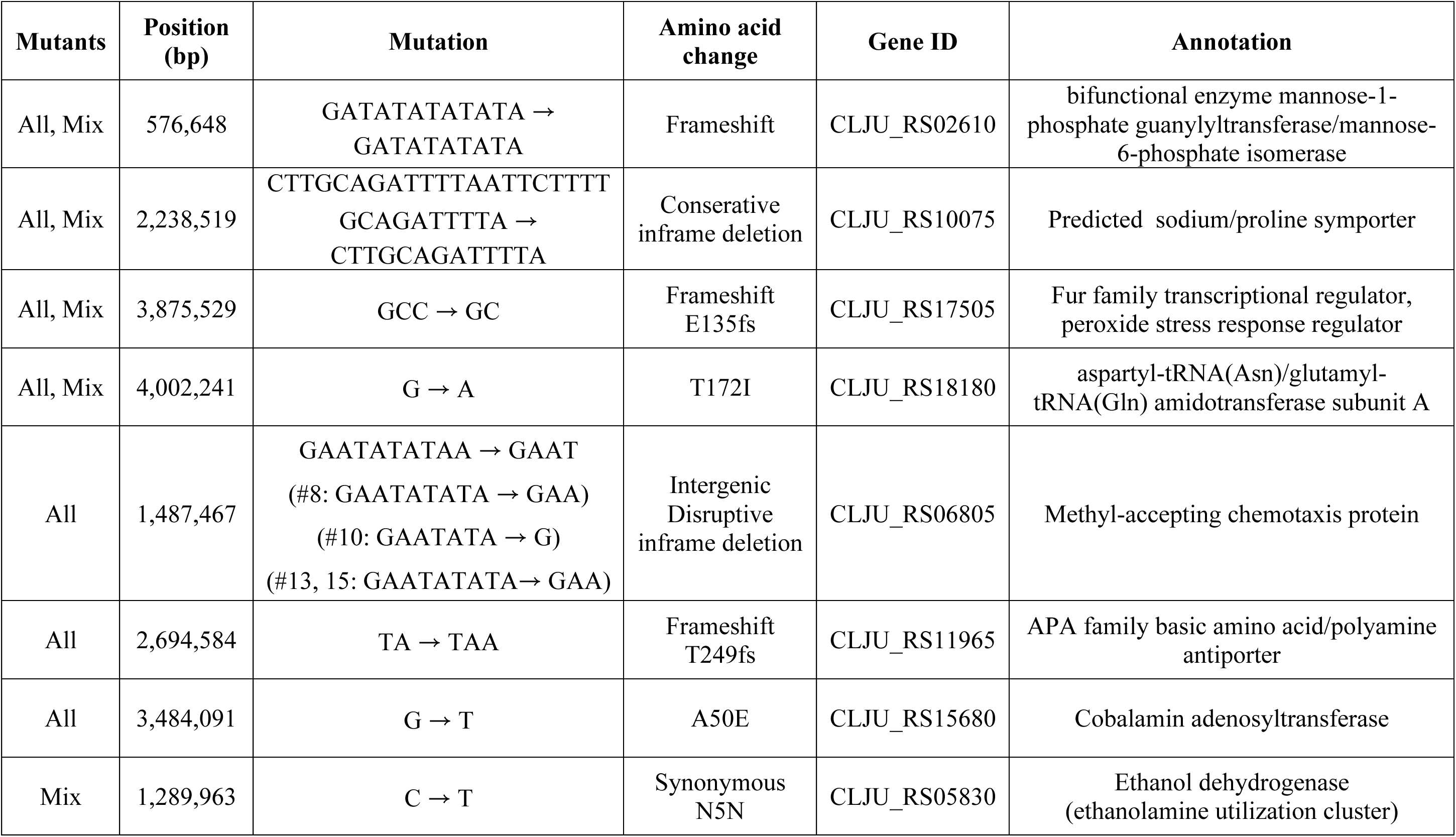
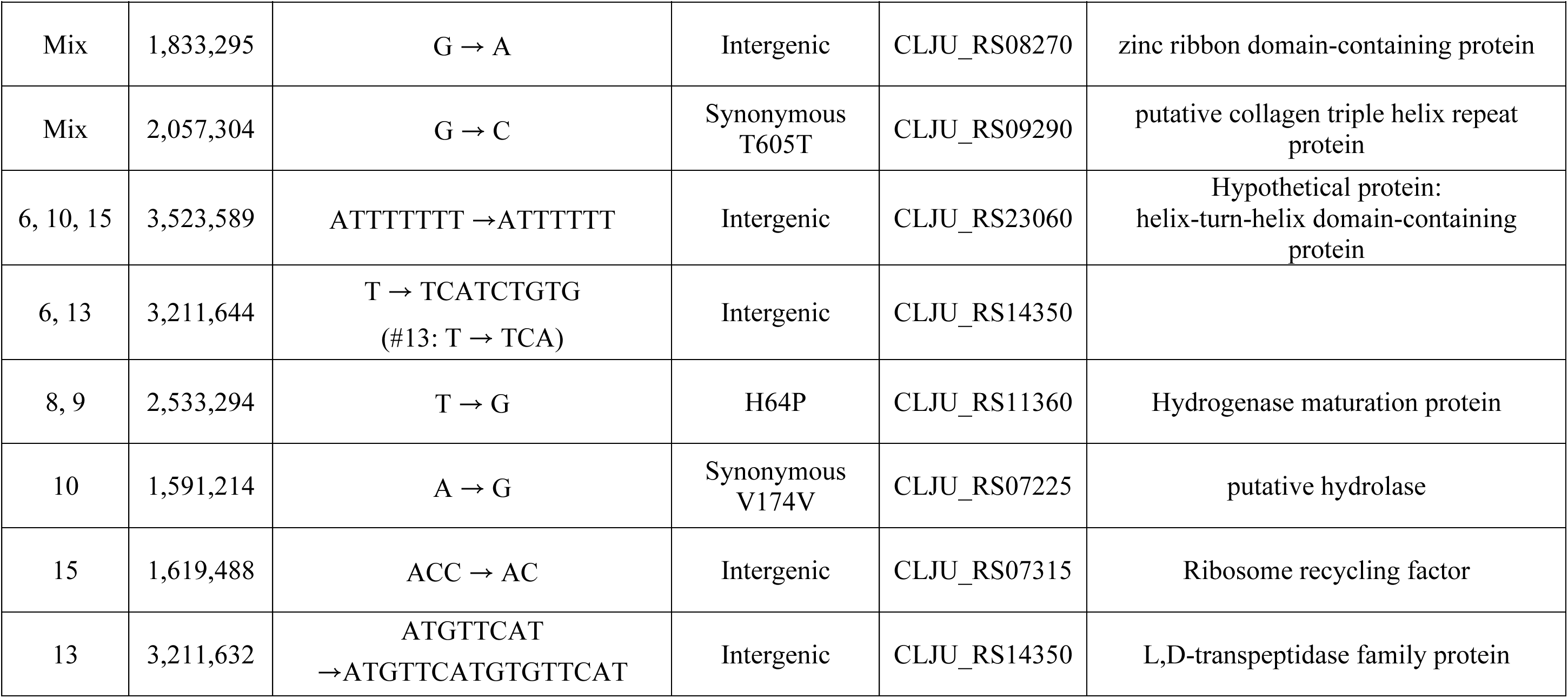
Mutations detected from isolated strains from the evolutionary adaptation experiments.

Compared to the reference genome, 22 mutations were identified by whole genome sequencing of the parental strain: 9 synonymous mutations, 6 mutations in intergenic regions, 6 missense, and 1 frameshift mutations. Among the 22 mutations, 3 mutations happened in glutamine and glutathione metabolism (CLJU_RS02190, CLJU_RS03040, and CLJU_RS10055), 12 mutations in genes encoding membrane proteins (CLJU_RS11595, CLJU_RS14955, CLJU_RS20060), 1 in gluconeogenesis (CLJU_RS14330), 3 in transcriptional regulators (CLJU_RS06385 and CLJU_RS07295), and 3 in genes encoding hypothetical proteins (CLJU_RS07330, CLJU_RS08235, and CLJU_RS22935).

Eight isolated strains were sequenced to analyze the mutations that had happened during evolutionary adaptation. The mixed population enriched directly in YTF medium from the iron granules was also sequenced to identify mutations that might have been missed during the isolation process (see Section 2.3). Seven mutations were found in all eight isolated strains (CLJU_RS02610, CLJU_RS06805, CLJU_RS10075, CLJU_RS11965, CLJU_RS15680, CLJU_RS17505, CLJU_RS18180) (Table 2). The genes for the bifunctional enzyme mannose-1-phosphate guanylyltransferase/mannose-6-phosphate isomerase (CLJU_RS02610), APA family basic amino acid/polyamine antiporter (CLJU_11965), and Fur family transcriptional regulator (CLJU_17505) were frameshifted. Inframe deletions were found in the genes encoding the methyl-accepting chemotaxis protein (CLJU_RS06805) and predicted sodium/proline symporter (CLJU_RS10075). Also, non-synonymous mutations were found in cobalamin adenosyltransferase (CLJU_RS15680) and aspartyl-tRNA/glutamyl-tRNA aminotransferase subunit A in all isolated strains.

Three mutations were found only from the mixed population (CLJU_RS05830, CLJU_RS08270, and CLJU_RS09290). Synonymous mutations were found in genes encoding ethanol dehydrogenase (CLJU_RS05830) and a putative collagen triple helix repeat protein (CLJU_RS09290). A mutation in an intergenic region for zinc ribbon domain-containing protein (CLJU_RS08270) was also found.

Six mutations were found in only small groups of isolated mutants (CLJU_RS07225, CLJU_RS07315, CLJU_RS11360, CLJU_RS14350, and CLJU_RS23060).

One of the mutations found in isolate #8 and #9 was the mutation on hypF (CLJU_RS11360) coding for a hydrogenase maturation protein. Among all the mutations, the mutation on hypF seemed to be the most directly relevant to iron utilization, based on literature study and the fact that hydrogenase is directly involved in hydrogen metabolism (8). Therefore, strain #8 was chosen for further tests in BES reactors.

### 3.6 Performance of an evolved mutant in BES reactors

In a BES reactor, wild-type *C. ljungdahlii* produced more metabolites (12.2 ± 0.9 mM acetate and 2.7 ± 0.4 mM formate) than isolate #8 (5.0 ± 0.1 mM acetate and 3.6 ± 0.4 mM formate) (Figure 6).

**Figure 6.**
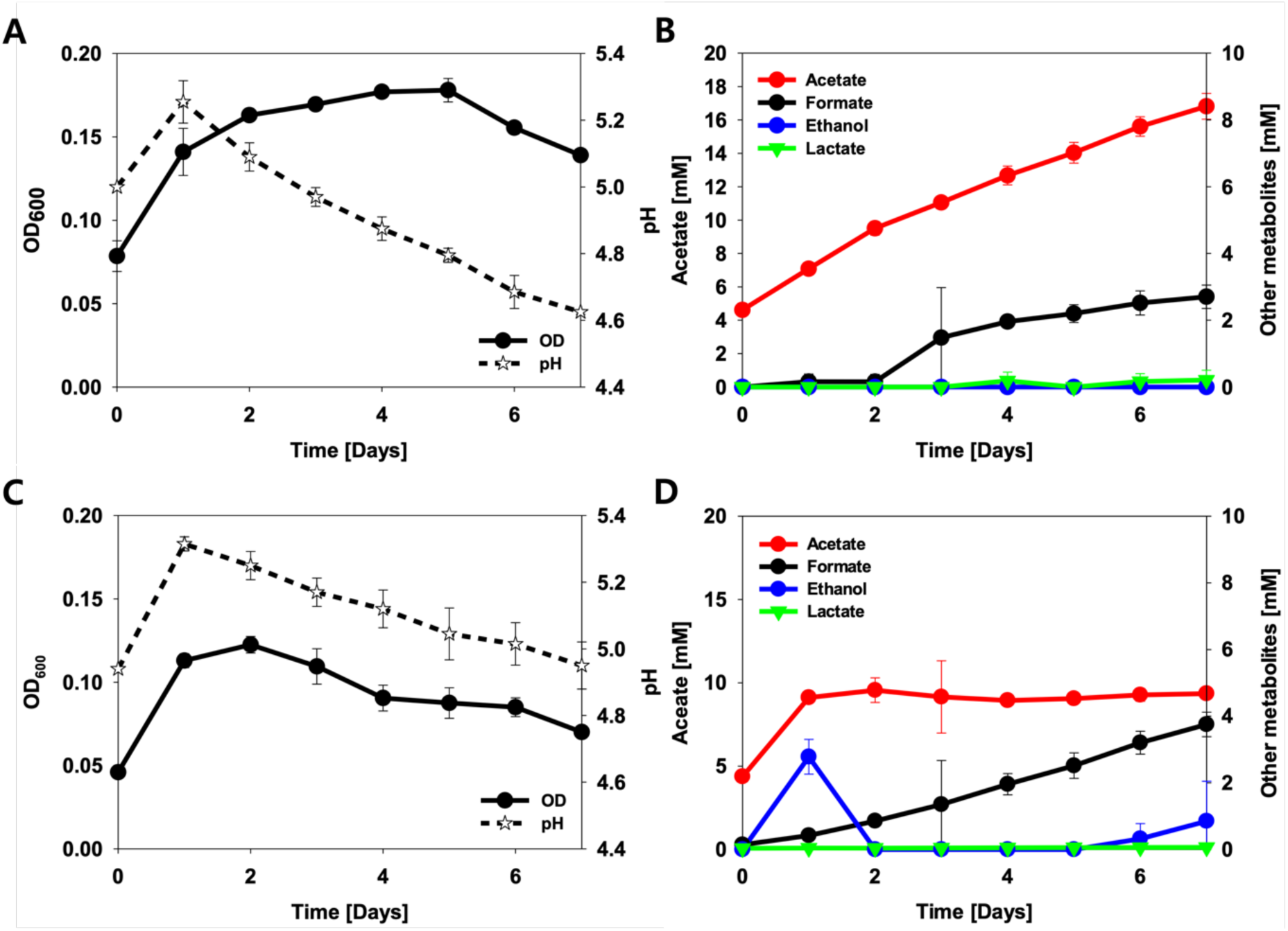
OD, pH and metabolite profiles of wild-type (A and B) and isolate #8 (C and D) in BES reactors. The error bars represent the standard deviations of the mean (n=2).

The OD of wild-type also increased to OD 0.177 ± 0.003 until day 4, while the maximum OD of isolate #8 was 0.122 ± 0.005 on day 2 (Figure 6 A and C). Interestingly, acetate production of isolate #8 was ceased after day 1, and formate became the main metabolite. Also, 2.8 ± 0.5 mM ethanol was detected on day 1 and re-assimilated on the next day, and ethanol production was detected from day 6 again.

## 4. Discussion

The aim of this study was to adapt *C. ljungdahlii* for better electron uptake from a solid donor, using iron as the sole electron source.

It was not possible to measure growth of *C. ljungdahlii* in the flasks for ALE due to unknown precipitations and accumulation of cells on the surface of the iron granules. Based on previous estimations of the doubling time of *C. ljungdahlii* in the same growth medium with a gas composition of [H_2_:CO_2_] = [80:20], which was reported as 34.5 ± 2.8 h (25), we estimate that the ALE experiment conducted for one year consisted of approximately 250 generations assuming planktonic cell growth under the H₂:CO₂ condition. Estimating enough time for desirable mutations to occur during the experiment was difficult, not only due to difficulties in estimating the growth rate but also due to difficulties in estimating to what extent a mutation is actually linked to an improved fitness (17). After the 32^nd^ medium replacement, the acetate production rates of the replicate cultures were increased by 255 % and 155%, respectively (Fig. 3). After three more medium replacements, the acetate production rate improved 10 and 6.5 times compared to the first three batches. It is difficult to argue that the specific acetate production rate of individual cells improved as we could not determine cell growth in this study. Still, as the acetate production rate increased radically during the 11^th^ and 12^th^ months of ALE, we assumed that major mutations had occurred during the experiment.

Selection pressure while isolating mutants is important when groups of diverse mutations are expected. As desirable mutations happening, the most fit strain becomes dominant during ALE. In this study, [H_2_:CO_2_] = [80:20] gas was used to enrich the population from the iron granules prior to isolation of individual colonies. Therefore, it is likely that the enriched mutants are the mutants growing faster under H_2_:CO_2_ condition, but not necessarily under CO_2_ and iron condition. Recently, a hypothesis that homoacetogen’s affinity to hydrogen plays an important role for utilization of insoluble electron donor was suggested (8). The hydrogen consumption characteristics of different acetogenic bacteria were compared and were hypothesized to relate to their capabilities of utilizing an insoluble electron donor (8). Therefore, enrichment of a population under high hydrogen partial pressure may have been inappropriate.

Screening desired mutations is critical for ALE experiments. We set to determine the heterotrophic and autotrophic growth of eight of the individual isolates. During the ALE on iron and CO_2_, it is plausible that evolution either favours cells with improved growth or cells capable of slowing down the metabolism to save available resources when the resources are limited (37, 38). The latter corresponds with our observations for autotrophic growth of isolates and acetate production rate on iron (Fig. 4 and 5).

Possibly, the mixed population of *C. ljungdahlii* after evolution on iron might have functioned synergistically under the given electron-and nutrient-limited condition, by dividing their roles in the population. In a mixed population, subpopulations can cross-feed on intermediate products, thereby reducing competition for limited resources. This may explain why the mixed population of *C. ljungahlii* showed improved performance after evolution on iron while each of the isolated mutants performed worse than the wild-type strain (Fig. 3 and 5).

After *Sporomusa ovata* was improved for increased BES performance through ALE on methanol, a mutation on *ackA*, acetate kinase, was assumed to be the major mutation that enhanced microbial electrosynthesis in a BES in the study (24). From the results, it can be hypothesized that before the ALE, the central metabolism of *S. ovata* had a lower capacity than the EEU of *S. ovata*. After adaptation, EEU was harmonized with the central metabolism to allow performance at the full capacity of the host strain in a BES. If *C. ljungdahlii*’s affinity to hydrogen plays an important role for EEU from a cathode, a mutation on hydrogenase or on other redox enzymes could be expected. A mutation on hydrogenase likely increases affinity to hydrogen but becomes slow as a trade-off, resulting in slow growth on hydrogen and on iron.

Mutations on other redox enzymes may also appear as a result of trying to find a new intra-cellular redox balance from low electron availability. The aim of this study was to evolve *C. ljungdahlii* strains on iron for improved extracellular electron uptake. However, no isolated mutants showed better acetate production under CO_2_ and iron condition compared to the parental strain (Fig. 5).

A mutation found in all the isolated mutants of *C. ljungdahlii* evolved on iron was in the gene coding for bifunctional mannose-1-phosphate guanylyltransferase/mannose-6-phosphate isomerase (CLJU_RS02610). This gene is connected to nucleotide sugar metabolism (39). This gene plays an important role in glycosylation, the main class of post-translational modifications. Because of the common mutation, this may be an important process during adaptation to iron (40). Methyl-accepting chemotaxis protein (CLJU_RS06805) is a receptor for bacterial chemotaxis (41). This receptor uses aspartate as an attractant (42). The mutation in gatA (aspartyl-tRNA/glutamyl-tRNA amidotransferase subunit A, CLJU_RS18180) might have been effected to enable the cells to find a better environment. If the mutation happened to improve chemotaxis, this could benefit the cells when there is a limited electron supply in the local medium (e.g. iron and cathode in a BES). Alternatively, decreased chemotaxis would be a way to save ATP.

The sodium/proline symporter (CLJU_RS10075) and the amino acid/polyamine symporter (CLJU_RS10075) were disrupted during ALE on iron. It is possible that the disruptions of the two transporters save cell resources required for the expression of membrane proteins while slowing down metabolism to match the EEU rate of *C. ljungdahlii*. CobO (Cobalamin adenosyltransferase,CLJU_RS15680) gene was also mutated (Table 4). Cobalamin (vitamin B12) is an important cofactor in Wood-Ljungdahl pathway, as well as in methionine synthase (43, 44). Recently, Dahle et al., reported methionine supplementation as a sulfur source is essential for *C. ljungdahlii* to grow in a defined medium (45). Therefore, it seems that *C. ljungdahlii* during evolutionary adaptation tried to improve supply of cobalamin as a cofactor for CO_2_ fixation and enhance methionine synthesis with a limited electron supply.

The omission of yeast extract in the medium, to promote the EEU from ZVI, might have challenged *C.ljungdahlii* growth under the given environment, resulting in the disruptions of the transporters for saving cell resources, cobalamin utilization for methionine synthesis, as well as methyl-accepting chemotaxis protein for saving ATP usage (46).

Another common mutation found in all isolates was located in CLJU_RS17505 (Fur family transcriptional regulator, peroxide stress response regulator). Its gene product appears to regulate Fe^2+^/Zn^2+^ uptake along with fur1, as well as expression of many other genes involved in metal-containing enzymes, including the metal ABC transporter of ATPase (CLJU_RS15665) and transcriptional regulator mntR (CLJU_RS16935) for metal transport. The mutation in CLJU_RS17505 may affect regulation of the uptake of excessive Fe^2+^ or Fe^3+^ present in the medium from ZVI oxidation.

The main aim of this study was to have mutations from *C. ljungdahlii* for increased electro-activity in a BES; based on the hypothesis that biological iron oxidation shares the same mechanism of biological cathode oxidation in a BES. The performance of isolate #8 was compared with wild-type in BES reactors. The cell growth and acetate production of isolate #8 did not increase after the first two days of growth. The initial growth observed during the first two days was likely due to the presence of yeast extract in the medium, as observed in a previous publication. (25), while the cell growth of wild-type was sustained until day 5 (Fig. 6). This suggest that the mutations during ALE did not happen as expected. There are three speculated reasons why ALE was not successful. 1) Limited nutrition of the defined medium was challenging for *C. ljungdahlii* to evolve for the better growth. This might be resolved by adding yeast extract since the contribution of yeast extract in cell growth is negligible without an electron donor, which was observed when *C. ljungdahlii* was cultivated under CO_2_ and yeast extract condition without any other electron source, yet yeast extract contains a lot of nutrients supporting cell growth such as amino acids, minerals, and vitamins (25). 2) Also, the limited time for each batch (maximum 2 weeks) due to pH increase made ALE without knowing the cell growth challenging as well as limited *C. ljungdahlii*’s growth pH. pH increase by iron oxidation was also a hurdle for ALE on iron using *C. ljungdahlii* (Fig. 1). Previously reported studies about isolation of acetogenic bacteria using CO_2_ + iron condition had to enrich the culture at least a month or two for each cultivation to get enough biomass (9, 11). Therefore, acetogenic bacteria that can grow on relatively high pH (pH 7.0 – 9.0), such as *Acetobacterium* spp. and *Sporomusa* spp., were isolated due to pH increase by iron oxidation (Fig. 1). pH control of flasks for ALE could make it possible longer ALE cultivations than without a pH control. The longer ALE cultivation would allow passage of a part of ALE culture to a sterile flask with iron granules, which enables selection of evolved strains with improved fitness (17). Lastly, 3) the selective pressure was not appropriate because the formed cells on iron granules kept in the same flask till the end without subculturing the part of evolved mutants in the new flask with fresh iron granules to make desired mutants dominant in the culture and accumulated. This approach also could have been challenged due to the limited cultivation time by pH increase.

Formate and ethanol are electron dense chemicals that can be used as soluble electron donor (47). While the cell chooses a strategy to slow down its metabolism under harsh growth condition, autotrophic bacteria also might need to convert gaseous substrate into soluble form that readily accessible and further metabolize in the future. That might be why *C. ljungdahlii* produces formate and ethanol as main metabolites after acetate production ceases (Fig. 6). Ethanol production (∼1.25mole ATP/mole ethanol) generates more ATP than acetate production (∼0.75 mole ATP/mole acetate) from CO_2_ and H_2_. Formate serves as a soluble electron storage instead of gaseous electron donor, H_2_, which can easily disappear (48, 49). Ethanolamine production from biofilm-forming *C. ljungdahlii* in a BES could explained in the same way (50). The same observation has been made from *Rhodobacter sphaeroides*, which is a polyhydroxyburate-forming photoautotrophic bacterium, in a BES (51).(52, 53). Biofilm-forming *R. sphaeroides* in a BES produced more polyhydroxybutyrate than planktonic cells (51). Therefore, it seems that autotrophic bacteria choose the survival strategy producing energy storage compound in an electron-limited condition (in a BES) and that *C. ljungdahlii* might have adapted to produce ethanol and formate as energy and electron storage compounds under CO_2_ and iron condition. The early cessation of cell growth of isolates #8 might be due to improved formate and ethanol production that uses reducing equivalents otherwise required for cell growth (Fig. 6).

This study is the first attempt on improvement of acetogenic extracellular electron uptake using an ALE method. *C. ljungdahlii* was evolved on iron as the sole electron source in order to improve its extracellular electron uptake. However, it seemed that the given conditions were too nutrient-limiting for *C. ljungdahlii* to adapt for better utilization of CO_2_ with Fe(0) as electron source. Whole genome seqencing results instead showed many mutations that can hypothetically be linked to nutrient uptake and energy utilization. An isolated mutant evolved to produce ethanol and formate after growth and acetate production stopped in a BES. It is speculated that *C. ljungdahlii* evolved to slow down its metabolism to match the rate of electron uptake and to store electrons and carbon in metabolizable metabolites when subjected to electron-limited conditions.

## Acknowledgement

This work was supported by the Swedish Energy Agency (grant number 46605-1) and the Area of Advance Energy, Chalmers University of Technology.

## Abbreviations

ALE: Adaptive laboratory evolution
BES: Bioelectrochemical system
EA: Evolutionary adaptation
EET: Extracellular electron transfer
EEU: Extracellular electron uptake
ZVI: Zero-valent iron

